# Fractal scaling of *C. elegans* behavior is shaped by insulin signaling

**DOI:** 10.1101/2021.12.02.471007

**Authors:** Yukinobu Arata, Itsuki Shiga, Yusaku Ikeda, Peter Jurica, Hiroshi Kimura, Ken Kiyono, Yasushi Sako

## Abstract

Fractal scaling governs the complex behavior of various animal species and, in humans, can be altered by neurodegenerative diseases and aging^1^. However, the mechanism underlying fractal scaling remains unknown. Here, we videorecorded *C. elegans* that had been cultured in a microfluidic device for 3 days and analyzed temporal patterns of *C. elegans* actions by fractal analyses. The residence-time distribution of *C. elegans* shared a common feature with those of human and mice^2–4^. Specifically, the residence-time power-law distribution of the active state changed to an exponential-like decline at a longer time scale, whereas this change did not occur in the inactive state. The exponential-like decline disappeared in starved *C. elegans* but was restored by culturing animals with glucose. The exponential-like decline similarly disappeared in insulin-signaling *daf-2* and *daf-16* mutants. Therefore, we conclude that insulin signaling regulates fractal scaling of *C. elegans* behavior. Our findings indicate that neurosensory modulation of *C. elegans* behavior by insulin signaling is achieved by regulation of fractal scaling. In humans, diabetes mellitus is associated with depression, bipolar disorder, and anxiety disorder^5^, which affect daily behavioral activities. We hypothesize that comorbid behavioral defects in patients with diabetes may be attributed to altered fractal scaling of human behavior.

## Main

In humans, ordinary daily activities^2^ and social behaviors, such as sports and communication^4^, tend to occur consecutively as a burst, and then suddenly cease for several days to months. These episodic bouts of behavior have also been observed in other vertebrates (mice^3^) and invertebrates *(C. elegans^6^,* flies^7^, and ants^8^). Activity time series of episodic behavioral bouts have non-periodic and intermittent patterns that appear repeatedly across a broad range of time scales. Such self-similar geometrical patterns across time scales are called fractal patterns; therefore, activity time series of animal behavior are characterized by fractal geometry. Neurodegenerative disorders (e.g., Alzheimer and Parkinson diseases) and aging^1^ have been shown to alter the fractal scaling of human behavior. These findings suggest that fractal scaling of animal behavior is regulated by neurophysiological mechanisms that are conserved among various animal species.

Daily and social-behavioral activities are affected by a broad range of neurophysiological states in the human brain. Among them, mood, an unconscious disposition to respond emotionally to objects or events encountered in life^5^, and a reward evaluation for each object or event^9^ are thought to play important roles. Insulin signaling has been shown to affect mood and the reward system in mouse and human brains^10^. Mice with a brain neuron-specific knockout of the insulin receptor gene (NIRKO mice) did not show defects in neuron proliferation or death during brain development; however, they did show age-related anxiety and depressive-like behaviors^11^. In humans, mood is improved by nasal administration of insulin in both healthy individuals and patients with diabetes, suggesting that insulin signaling is involved in mood control^12^. Insulin signaling, which has evolved in relation to the mood and reward systems in brain in higher animals, modulates the relation between olfactory stimuli and behavior in nematodes and flies^10^. Thus, insulin signaling is an evolutionarily conserved signaling system that coordinates external stimulation and animal behavior. However, how insulin signaling affects the fractal scaling of animal behavior remains uninvestigated.

In the present study, we applied a genetic analysis of the fractal scaling of animal behavior by studying *C. elegans* behavior. Alternative switching between an actively moving state (“active state”) and an inactive state in episodic behavior is a common feature of various animal species. Therefore, we dissected the fractal scaling of *C. elegans* behavior on a two-state transition model^6^. Generally, kinetics that governs the state transition can be inferred from statistical properties, such as the frequency distribution and temporal correlation of experimentally measured residence times in each state. Inferred kinetics provides insights into the underlying mechanisms that drive the state transition. Through longitudinal videorecording of *C. elegans* swimming behavior, we found that state transitions between active and inactive states in *C. elegans* episodic behavior are driven by kinetics that determines residence times by following frequency distributions and temporal correlations with fractal properties. Therefore, we refer to the kinetics as “fractal kinetics”^6^.

Next, we extended the function of the microfluidic device for culturing *C. elegans* with food bacteria. Our observations revealed that the fractal kinetics of *C. elegans* behavior is regulated by insulin signaling. Based on recent neuronal network modelling and molecular biological studies, we discuss the possibility that insulin signaling regulates neural activity in the brain to modulate fractal scaling of *C. elegans* behavior. We also discuss the applicability of this mechanism for mood disorders that are comorbid with diabetes mellitus in humans. We propose that fractal behavioral analysis can provide a more integrated clinical view of psychiatric symptoms in patients with diabetes, which may contribute to the development of new diagnostic indices and improvement of clinical treatment.

### Residence-time power-law distributions in active and inactive states in *C. elegans* episodic behavior

To study the effects of diet on fractal scaling of *C. elegans* behavior, we constructed a new microfluidic device composed of an array of 50 chambers for culturing individual animals by perfusing M9 buffer containing food bacteria (WormFloII, Fig. 1). We recorded *C. elegans* swimming under controlled chemical, temperature, and light intensity conditions at 20 frames per second (fps) for 3 days^6^. By analyzing recorded movies using an image-processing program^6^, we obtained time series of behavioral activity with 10^5^ time points (Fig. 2a-e). In the activity time series, we confirmed that fed wild-type animals cultured on the device showed repeated active and inactive episodes (Fig. 2b, c), as observed in *C. elegans* cultured in liquid and solid agar medium^6^. These findings indicate that our culture system allowed us to observe the physiological behavior of *C. elegans.*

**Fig. 1:**
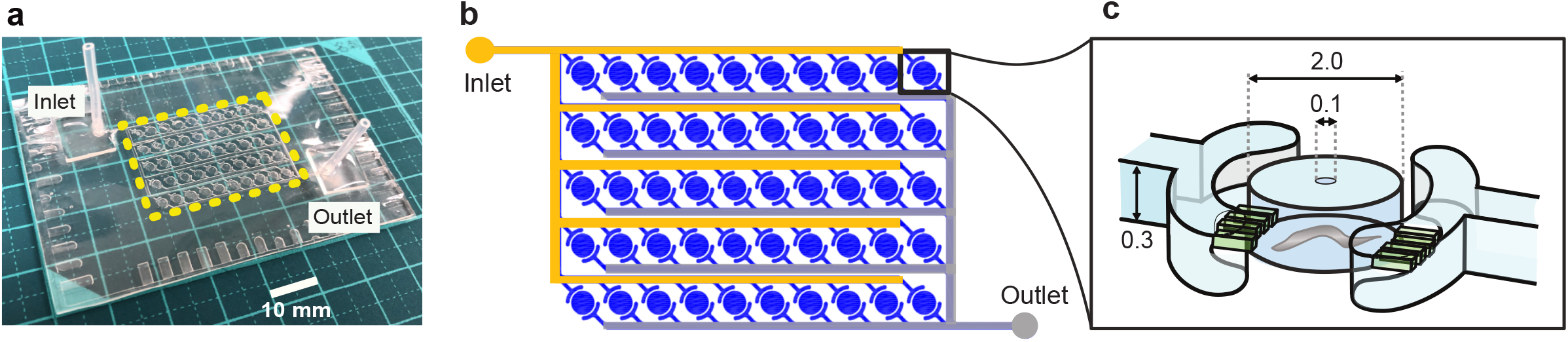
WormFloII microfluidic device for culturing *C. elegans* with food bacteria in biochemical isolation. **a,** WormFloII photo. Inlet and outlet portals for liquid media are shown. Yellow dashed line outlines 50 chambers for individually culturing *C. elegans.* **b,** WormFloII schematic. Each chamber is directly connected to supply (orange) and drain (gray) channels to achieve biochemically independent environments. **c,** Chamber schematic. Chambers are caged with junctional micro-slit channels (50-μm width and height). Animals are introduced from 0.1-mm hole at chamber roof. Hole is shielded with PDMS sheet before supplying liquid media.

**Fig. 2:**
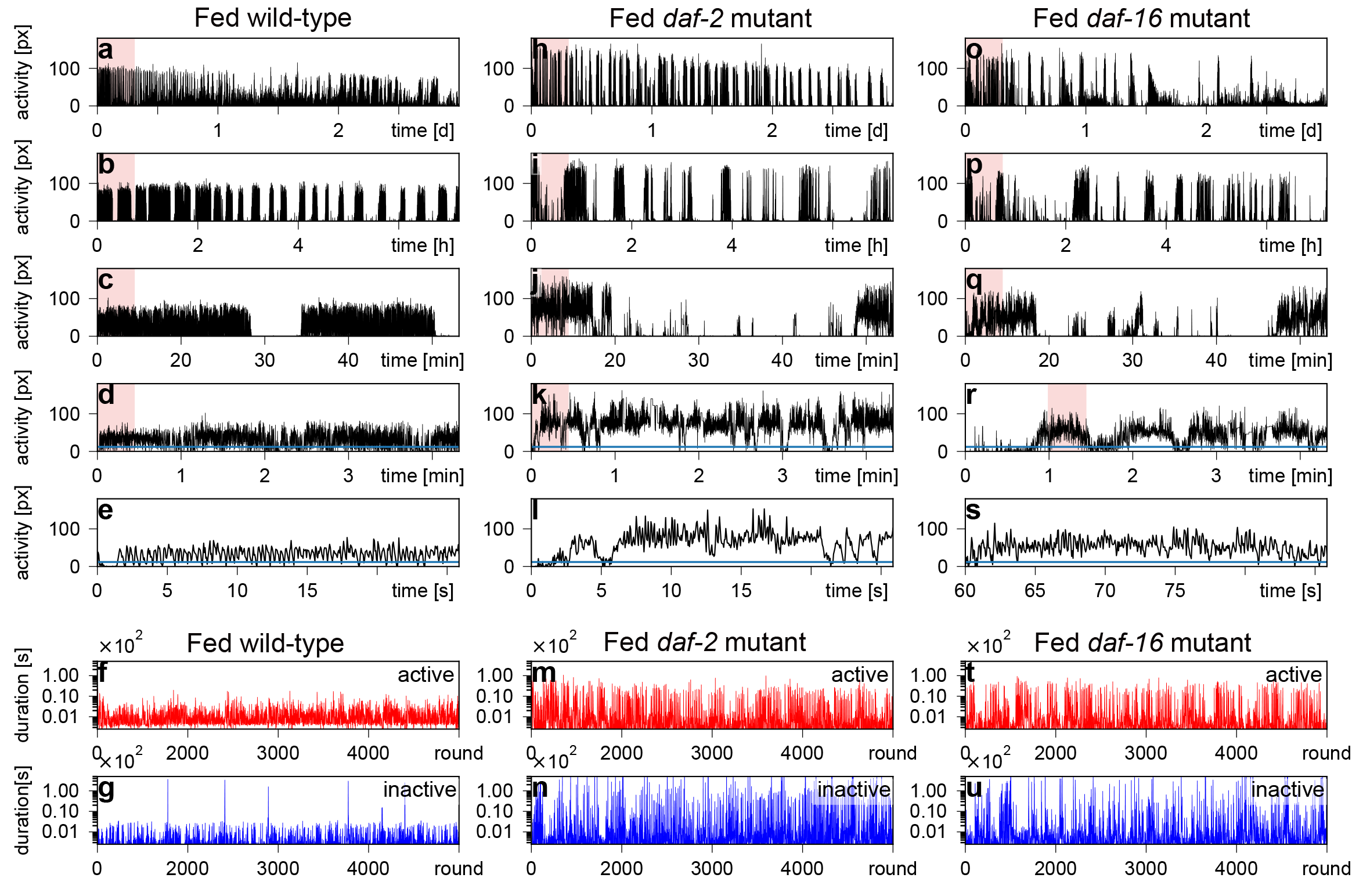
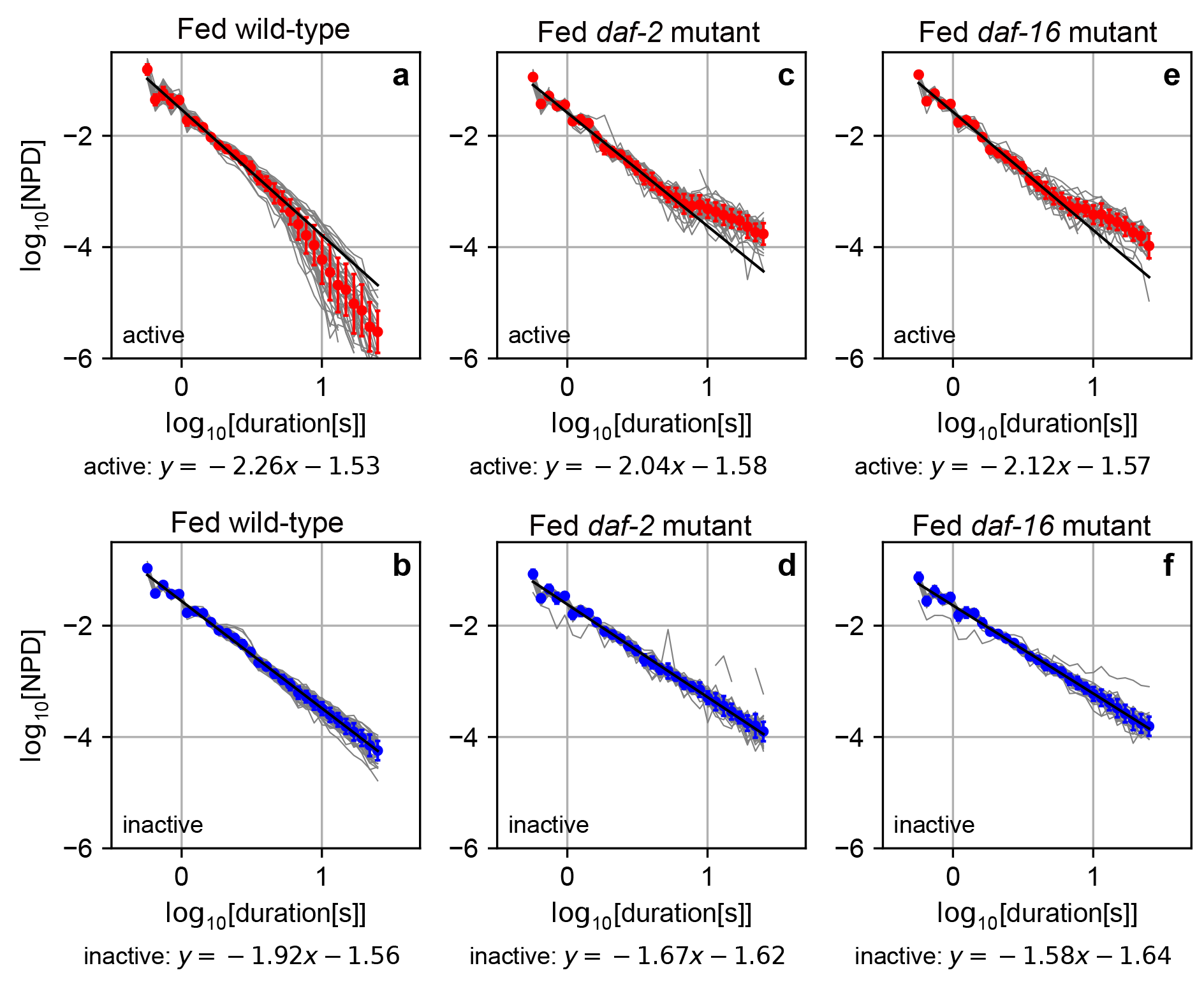
Activity time series of episodic behavior of fed *C. elegans*. Swimming activity time series of wild-type (**a-e**), *daf-2* (**h-l**), and *daf-16* (**o-s**) fed animals for 3 days. Red-marked regions are magnified in graphs immediately below (e.g., all of **b** represents red region from **a**; all of **c** represents red region from **b**; etc.). DRSs for active (red) and inactive state (blue) in wild-type (**f**, **g**), *daf-2* (**m**, **n**), and *daf-16* (**t**, **u**) fed animals were obtained from activity time series for 3 days (**a**, **h**, **o**). px: number of pixels where animals moved from previous frame.

To analyze the fractal scaling of *C. elegans* behavior based on a two-state transition model, we measured the residence times of the active and inactive states, which alternatively appeared along the activity time series. Residence-time series in the active or inactive state were plotted across the round as “duration round series” (DRS) (Fig. 2f, g), analogous to “activity time series”. DRS in fed wild-type animals revealed that residence times in the active state varied from sub-seconds to 10 seconds, whereas residence times in the inactive state varied from sub-seconds to 100 seconds (Fig. 2f, g). In the inactive state, residence times followed a power-law distribution in the range of sub-seconds to >10 seconds (Fig. 3b). In the active state, residence times followed a power-law distribution in a shorter range, from sub-seconds to <10 seconds (Fig. 3a). This power-law distribution indicates that the appearance frequency of the residence time decreased on the time scale in a certain ratio across a broad range of residence times. In other words, the appearance frequency decreases in a self-similar manner, which is indicative of fractal scaling in the residence time. Interestingly, at a longer time scale, the frequency distribution of residence time in the active state showed a faster decline than the power-law distribution. Such a convex decline in the log-log plot is seen in an exponential distribution. A similar combination of frequency distributions with and without the exponential-like decline in active and inactive states, respectively, was reported in Japanese quail, mice, and humans^2–4,13^. Thus, the frequency distribution of *C. elegans* episodic swimming has a common scaling property to vertebrates.

Next, we studied the behavioral activity of *C. elegans* that had been cultured in M9 buffer alone (starved wild-type animals) or cultured with 1 g/L glucose (glucose-fed wild-type animals) (Extended Data Fig. 1). The residence-time distribution of starved wild-type animals did not show a detectable exponential-like decline in either the active or inactive state (Extended Data Fig. 2a,b)^6^. In glucose-fed wild-type animals, the exponential-like decline was restored in the active state (Extended Data Fig. 2c, d), raising the possibility that insulin signaling is involved in regulation of fractal scaling of *C. elegans* behavior. To test this possibility, we studied the *daf-2* (Fig. 2h-n) and *daf-16* (Fig. 2t-u) insulin-signaling mutant animals. *daf-2* and *daf-16* are mutants of the insulin receptor gene and of the downstream forkhead transcription factor gene, respectively^14^. In both cases, the insulin-signaling mutants showed increased frequency of the long-lasting active state compared to wild-type animals, such that the exponential-like decline at the longer time scale disappeared in the active state (Fig. 3c-3f). Additionally, through quantitative analysis of the power-law distribution, we found that the absolute value of the power-law exponent in the active state at the shorter time scale became larger in an insulin signaling-dependent manner (*p* < 0.05, Extended Data Fig. 3 and Supplementary Discussion). Therefore, we conclude that the mechanism to determine residence-time distribution in the active but not the inactive state is controlled by insulin signaling (Supplementary Discussion).

### Long-range correlation in duration-round series of active and inactive states in C. *elegans* episodic behaviors

To further study the insulin signaling-dependent control of fractal scaling of *C. elegans* behavior, we focused on the autocorrelation of DRSs. When the autocorrelation of one-dimensional data series declines with time lag *τ* in a power-law manner (*C*(*τ*) ~ *τ~^γ^*), such an autocorrelation is referred to as “long-range correlation,” due to the long tail in the power-law distribution. A power-law distribution of autocorrelation indicates that autocorrelation declines in a certain ratio across a broad range of time-lags, i.e., autocorrelation declines in a self-similar manner, which is indicative of fractal scaling across the round of residence times.

To study the long-range correlation in fractal scaling of *C. elegans* behavior, we employed higher-order detrending moving-average analysis (DMA)^15^. In DMA and its two-variable extension, detrending moving-average cross-correlation analysis (DMCA), when the fluctuation functions (*F*(*s*) or *F*^(1,2)^(*s*), equations (1), (2)) follow a power law with scale (*s*) (*F*(*s*)~*s^α^* or *F*^(1,2)^(*s*)~*s^α^*), *α* corresponds to the Hurst exponent (*H*)^15,16^. *H* obtained by DMA and DMCA has a direct mathematical link with other conventional indices for long-range correlation: i.e., the scaling exponent *γ* in autocorrelation (*γ* = 2 — 2*α*, for 0 < *γ* < 1) and the scaling exponent *β* in power spectral density *P*(*f*)~*f^β^*, where *f* is the frequency (*β* = 2*α* – 1, for *β* > −1)^17^. Compared to conventional algorithms, DMA has several advantages for estimating long-range correlation for scaling exponent *γ* or *β*, due to the availability of a fast algorithm and improved trend removal process^15^. When *H* = 0.5, the time series had no temporal correlation (i.e., like white noise), whereas when 0.5 < *H* < 1, the time series had a long-range correlation. Although long-range correlation of the time series cannot be simply extended to *H* > 1 due to 0 < *γ* < 1, fractal scaling of the time series can be characterized by *H* > 1. When *H* is larger (*H* > 0.5), there is a stronger tendency for values in the time series to continuously increase or decrease^6,18^. Therefore, in our study, we classified fractal scaling of the time series as “no memory” at *H* = 0.5, “weak fractal memory” for 0.5 < *H* < 1, and “strong fractal memory” at *H* > 1. Note that we used *H* of the integrated time series to characterize fractal scaling of the original time series, by following a standard algorithm of DMA (Methods).

The Hurst exponent of DRS of the active state (active DRS) at shorter round scale (< 100 rounds, *H*_*a*1_ = 0.70) and that at longer round scale (> 100 rounds, *H*_*a*2_ = 0.72) (Fig. 4a), and the Hurst exponent of DRS of the inactive state (inactive DRS)(*H_i_* = 0.68, Fig.4a) indicate that the active and inactive DRS have weak fractal memories, consistent with our previous study^6^. We did not find strong evidence for insulin signaling-dependent control of the mechanism to determine temporal correlations in active and inactive DRSs (Extended Data Figs. 4, 5a-f and Supplementary Discussion).

**Fig. 4:**
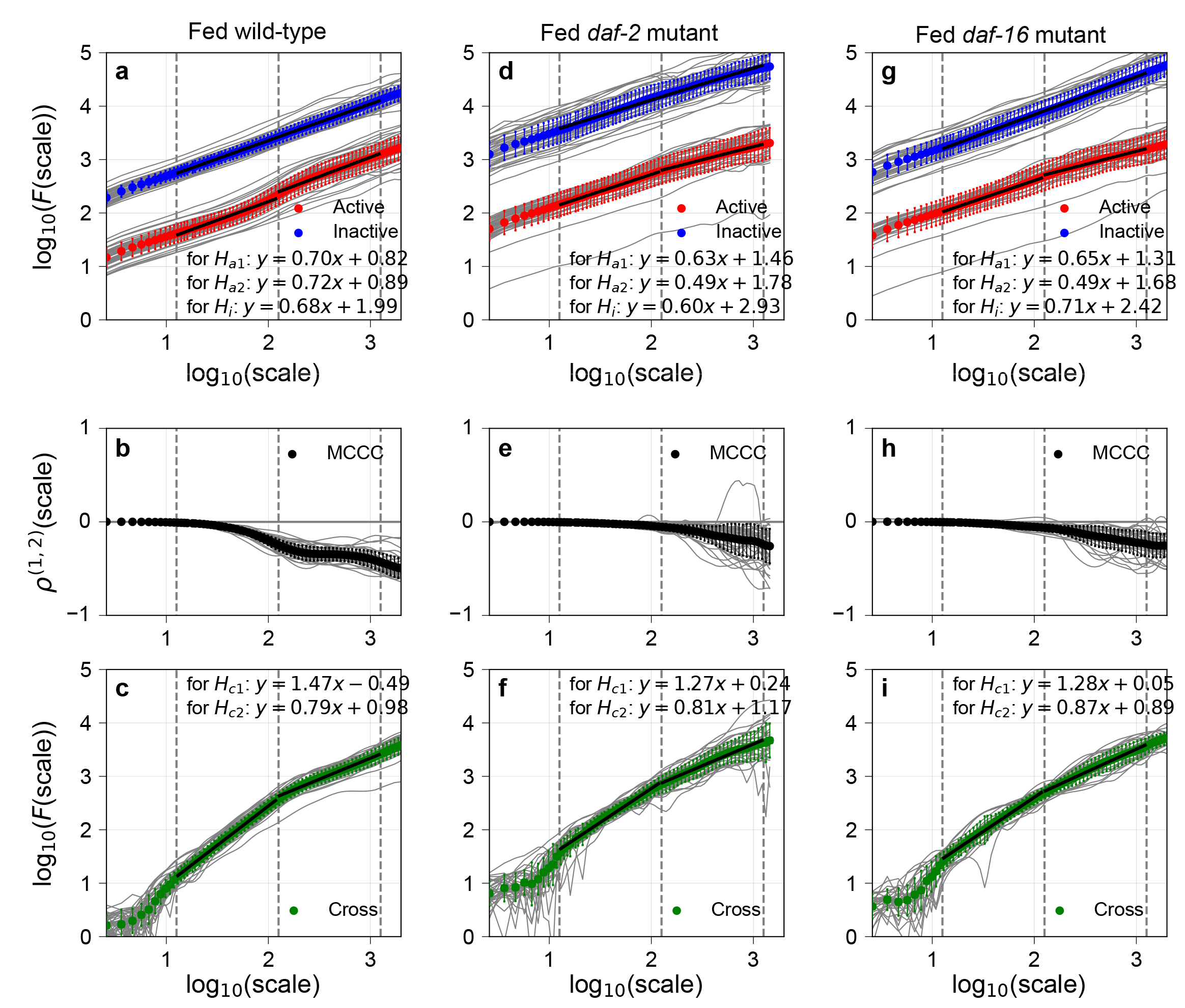
Long-range auto-/cross-correlations and multiscale cross-correlation coefficients in fed *C. elegans* behavior. Averaged noise function *F*(*s*) of active (red) and averaged cross-noise function *F*^(1,2)^(*s*) (green) among individual animals (grey) were fit with linear function from 1.1 to 2.1 and from 2.1 to 3.1 across scale (*s*) at shorter and longer round scale, respectively. Averaged noise function *F*(*s*) of inactive (blue) DRSs was fit from 1.1 to 3.1. Averaged multiscale cross-correlation coefficient (MCCC; black) among individual animals (grey) were plotted against scale (*s*) for fed wild-type (**b**), *daf-2* (**e**), and *daf-16* (**h**). Error bars are standard deviation.

**Fig. 5:**
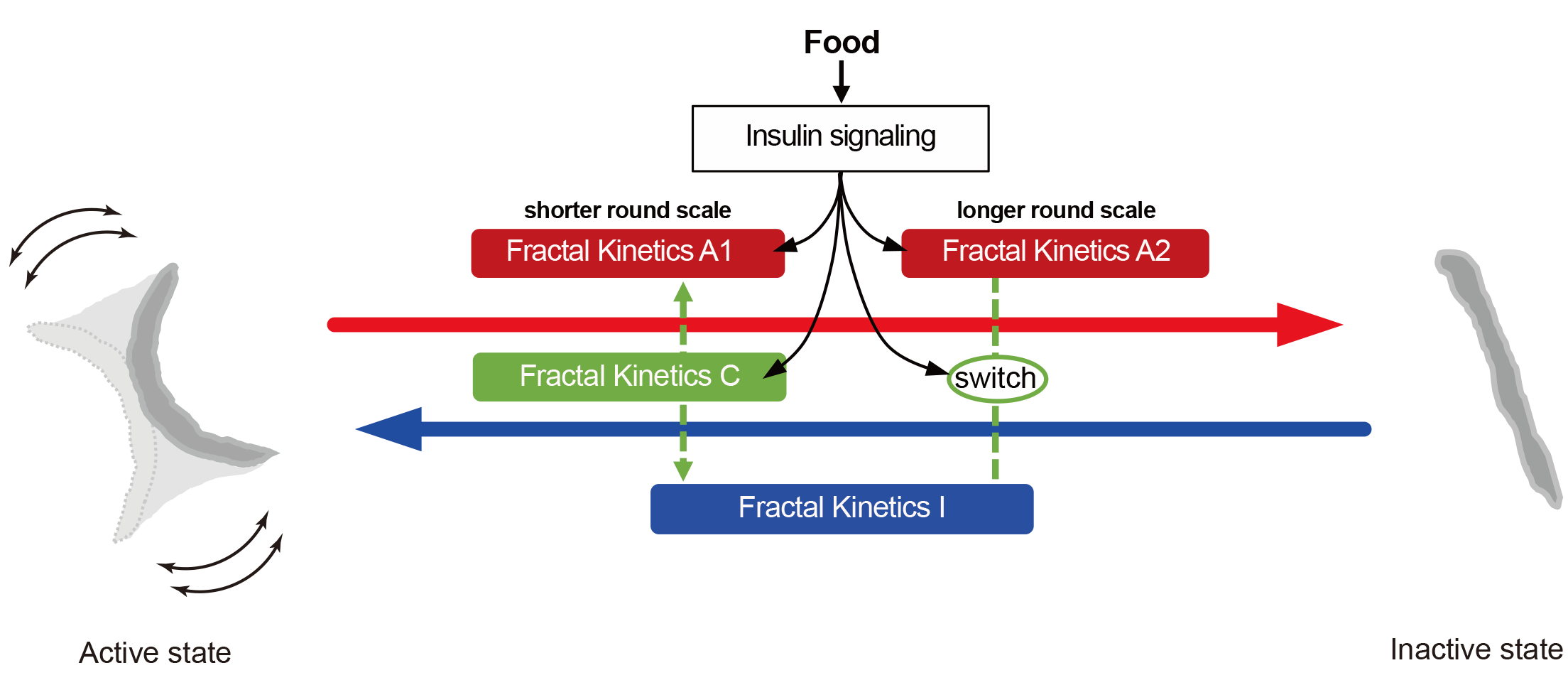
Two-state behavioral transition model. State transition from the active to inactive state is driven by Fractal Kinetics A1 at shorter round scale (which is affected by Fractal Kinetics C) and is driven by fractal Kinetics A2 at longer round scale (which is affected by Fractal Kinetics I). State transition from the inactive to active state is driven by Fractal Kinetics I, which is affected by Fractal Kinetics C at the shorter round scale, and is affected by Fractal Kinetics A2 at the longer round scale. Insulin signaling targets Fractal Kinetics A1, A2, C, and the lateral linking mechanism.

### Cross-correlation between duration round series of active and inactive states in *C. elegans* episodic behavior

To study the relation between active and inactive DRSs, we estimated the cross-correlation coefficients between two DRSs at various temporal scales (multiscale cross-correlation coefficient, *ρ*^(1,2)^ (*s*), equation (3)). Fed wild-type animals showed a remarkable negative correlation at longer round scales (Fig. 4b, Extended Data Fig. 6). The negative correlation at longer round scale (*log*_10_(*s*) = 2.5) in fed wild-type animals (*ρ*_2.5_ = −0.35) was significantly weakened in insulin-signaling mutant animals (*ρ*_2.5_ = −0.10 and *ρ*_2.5_ = −0.10 in fed *daf-2* and fed *daf-16* mutants, respectively, Fig. 4e, h, *p* < 0.05, Extended Data Fig. 6) and in starved wild-type animals (*ρ*_2.5_ = —0.09, Extended Data Fig. 4b, and *p* < 0.05, Extended Data Fig. 6). The negative correlation in starved wild-type animals (*ρ*_2.5_ = −0.09) was restored in glucose-fed wild-type animals (*ρ*_2.5_ = −0.13, Extended Data Fig. 4e, *p* < 0.05, Extended Data Fig. 6). These results indicate that there is a lateral linking mechanism between the two fractal kinetics (to determine active and inactive DRSs) at longer round scale, whose switch is modulated by insulin signaling.

Next, to study the *long-range* cross-correlation between active and inactive DRSs, we employed DMCA. In fed wild-type animals, Hurst exponents of a cross-correlated component between active and inactive DRSs at a shorter round scale (*H*_*c*l_ = 1.47) and at a longer round scale (*H*_*c*2_ = 0.79) (Fig. 4c) indicate that active and inactive DRSs contain a cross-correlated fractal component with strong fractal memory at a shorter round scale and weak fractal memory at a longer round scale. At a shorter round scale, *H*_*c*1_ in fed wild-type animals (1.47) was decreased in insulin-signaling mutants (1.27 and 1.28 in fed *daf-2* and fed *daf-16* mutants, respectively; Fig. 4f, i, *p* < 0.05, Extended Data Fig. 5g, h) and in starved wild-type animals (1.00) (*p* < 0.05, Extended Data Fig. 5g, h). Additionally, *H*_*c*1_ in starved wild-type animals (1.00) was restored in glucose-fed wild-type animals (1.20) (p < 0.05, Extended Data Fig. 5g, h). These results indicate that the strength of fractal memory in the cross-correlated component, unlike the fractal memories of active and inactive DRSs, is controlled by insulin signaling.

To our knowledge, there is no simple model to increase the strength of fractal memory by coupling simple models that generate time series with a weaker fractal memory^19^. Therefore, we consider that DRS with strong fractal memory generated by an upstream fractal kinetics is provided to both the active and inactive DRSs as a pseudo-cross-correlated component via a vertical interaction mechanism. On the other hand, at longer round scale, *H*_*c*2_ in fed wild-type animals (0.79, Fig. 4c) was comparable to the fractal memory of the active or inactive DRSs in fed wild-type animals (0.72 and 0.68, Fig. 4a). These values did change significantly in insulin-signaling mutants (0.81 and 0.87 in fed *daf-2* and fed *daf-16* mutants, respectively; Fig. 4f, i, Extended Data Fig. 5i, j). Due to the comparable strength of fractal memory in *H*_*c*2_ compared to those in *H*_*a*2_ and *H_i_*, the presence of an upstream fractal kinetics for a cross-correlated component remained unclear. It is possible that the DRS generated by fractal kinetics to generate active or inactive DRS was provided to the other DRS as a cross-correlated component via a lateral linking mechanism, which may be the same as the lateral linking mechanism found by the multiscale cross-correlation coefficient above. How insulin signaling-dependent behavioral control detected by DRS-based analyses alter the temporal activity patterns of *C. elegans* behavior are discussed in the Supplementary Discussion (Extended Data Figs. 7, 8).

## Discussion

### Fractal scaling of *C. elegans* behavior and insulin signaling-dependent control of neural activity in brain

Based on our fractal analyses, we dissected the fractal scaling of *C. elegans* behavior using a two-state transition model between active and inactive states (Fig. 5). State transition from the active to inactive state is driven by kinetics that determine residence time in the active state by following power-law and exponential-like distributions at shorter and longer time scales (Fig. 3a). The temporal correlation of residence times across the round is determined by following weak fractal memories with distinct strengths at shorter and longer round scales (*H*_*a*1_ and *H*_*a*2_; Fig. 4a). We refer to such kinetics as Fractal Kinetics A1 and A2, respectively. The temporal correlation determined by Fractal Kinetics A1 is affected by Fractal Kinetics C, which determines the temporal correlation by following strong fractal memory, via the vertical linking mechanism (*H*_*c*1_; Fig. 4c). The temporal correlation determined by Fractal Kinetics A2 is affected by Fractal Kinetics I via the lateral linking mechanism (*ρ*_2.5_; Fig. 4). On the other hand, the state transition from the inactive to active state is driven by kinetics that determines the residence time in the inactive state by following power-law distributions (Fig. 3b). The temporal correlation of residence times across the round is determined by following weak fractal memory (*H_i_*,; Fig. 4a). We refer to such kinetics as Fractal Kinetics I. The temporal correlation determined by Fractal Kinetics I is affected by Fractal Kinetics C via the vertical linking mechanism at shorter round scale (*H*_*c*1_; Fig. 4c) and by Fractal Kinetics A2 via the lateral linking mechanism at longer round scale (*ρ*_2.5_; Fig. 4). Insulin signaling modulates the mechanism for determining the residence-time distribution in Fractal Kinetics A1 and A2 (Fig. 3), and the mechanism for determining fractal memory in Fractal Kinetics C (*H*_*c*1_; Fig. 4c, f, i). Insulin signaling also targets the switch for the lateral linking mechanism between Fractal Kinetics A2 and I (*ρ*_2.5_; Fig. 4b, e, h) to shape fractal scaling of *C. elegans* behavior.

The generator of fractal kinetics may reside in the neural network in *C. elegans* brains. Power-law distributions have been observed in the duration of the sequential firing of neurons, called a “neuronal avalanche”, on cultured rat brain slices^20^ and in cat, monkey, and human brains *in vitro* and *in vivo*^21^. The power-law exponent of neuronal avalanche dynamics in the brain varies depending on the resting or task-performing behavioral state^22^ and on a wide range of neurogenic or psychiatric diseases in humans^23^. Exponents of the power-law distribution of neuronal avalanche are commonly distributed around −2, which approximately coincides with the power-law exponents determined by behavioral fractal kinetics A1 and A2 in *C. elegans* (Extended Data Fig. 3). A power law with a slope of −2 (second power law) in behavioral fractal kinetics is widely observed in invertebrates and vertebrates, including *Drosophila*^7^, Japanese quail^13^, mouse^3^, and human^2,4^. These observations strongly suggest that the power-law distribution in brain neuronal avalanche underlies the power-law distribution in behavioral fractal kinetics. Moreover, the second power law of neuronal avalanche has been reproduced by various theoretical neural network (NN) models^21^, including: stochastic NN models based on second-order phase transition (referred to as “criticality”)^24^, a deterministic model based on a feed forward-type NN model^25,26^, and another deterministic model based on the “edge of chaos” model^27^. Together, these findings suggest that behavioral fractal kinetics are derived from the conserved property of collective neural dynamics in animal brains among a wide variety of species.

Previous theoretical model analyses and subsequent experiments consistently explain the altered power-law residence-time distribution in fractal kinetics A1/A2 in insulin-signaling mutants (Fig. 3c, e). In previous stochastic and deterministic NN models, the power-law distribution of neuronal avalanche duration changes to an exponential-like decline at longer time scale, as we found in fed wild-type animals (Fig. 2a), when the maximum number of neurons in the model is reduced or neural connectivity is weakened^27–29^. These models suggest that negative interventions on propagation of neuronal firing in the network cause the exponential-like decline. This prediction has been validated in experiments using cultured rat brain slices. When a brain slice was cultured with an inhibitory neuron antagonist (i.e., picrotoxin), the average neuronal avalanche duration was elongated, such that the exponential-like decline disappeared from the frequency distribution observed in a brain slice cultured without the antagonist. With an inhibitory neuron antagonist, the frequency distribution at the longer time scale was beyond the power-law distribution^20,30^, similar to what we observed in fed *daf-2* and fed *daf-16* animals (Fig. 3c, e). These theoretical and experimental evidences suggest that the exponential-like decline in the residence-time distribution observed in *C. elegans* and other animals^3,4,13,31^ may be derived from some negative effects on brain neural activity by insulin signaling.

Previous molecular biological analyses have shown that insulin signaling has multifaced functions in brain^32,33^. In mammals, insulin acts as a neuropeptide to activate the GABA inhibitory ganglia in amygdala^34^, a key brain region connecting emotion/mood with food intake, providing a negative effect on neural activity. In *C. elegans,* GABA inhibitory D type-motor neurons (D-MNs) are involved in behavioral threat-reward decision making^35^. Our findings, together with previous theoretical and experimental studies, raise the possibility that insulin signaling in *C. elegans* activates GABA inhibitory neurons and alters the power-law neuronal avalanche distribution to be exponential-like, thereby changing the accompanying power-law residence-time distribution to be exponential-like.

How fractal kinetics I and C are generated, and how their interactions in the kinetic regulatory pathway (Fig. 5) are achieved in the *C. elegans* brain, remain unknown. We assume that the mechanism to determine the power law and exponential-like distribution of residence time in the active state shared in fractal kinetics A1 and A2 is generated from the specific neuronal network containing D-MNs in *C. elegans* (or the neuronal network in mammalian amygdala). In that case, fractal kinetics I and C may be generated from other distinct functional units in *C. elegans* and mammalian brains, and these functional units for fractal kinetics may structurally interact with each other in brain, as shown in Fig. 5. Considering the universality of power-law exponents of animal behavior and neuronal avalanche, the consistency of model prediction and experiments, and the evolutionary conservation of insulin signaling, the insulin-dependent control of fractal kinetics in the kinetic regulatory pathway (Fig. 5) may be conserved in fractal behavioral scaling in other animals.

### Insulin signaling and fractal human behavior

In humans, diabetic mellitus is associated with mood disorders, such as depression, bipolar disorder, and generalized anxiety disorder^5^, which affect daily behavioral activities, including food intake, sleep, communication, or social activities. These activities occur at different time scales. Our *C. elegans* fractal behavioral analysis raises the possibility that daily behavioral disorders in patients with diabetes at different time scales may be attributed to a disorder in fractal scaling of human behavior. This possibility could be tested through long-term measurements of human behavior in patients with diabetes and their evaluation by statistical fractal indices determined by the power-law residence-time distribution, cross-correlation coefficient (*ρ*^(1,2)^(*s*)), and long-range cross-correlation (*H*_*c*1_) (which, in the current study, were found to be regulated by insulin signaling). In parallel, statistical fractal indices obtained from behavioral dynamics in healthy individuals and patients with diabetes can be evaluated by a theoretical NN model representing the human brain structure^36^. Model analyses would provide additional multifaceted connections of multiple properties in behavioral fractal kinetics with brain neural activity. Together, the combination of long-term measurements, fractal statistical analysis, and theoretical neurodynamic modeling of fractal scaling of human behavior is expected to provide a more integrated clinical view of psychiatric symptoms in human patients with diabetes, which could contribute to the development of new diagnostic indices and the improvement of clinical treatment.

## Methods

### *C. elegans* strains and maintenance

*C. elegans* strains Bristol N2 (wild-type), CB1370 *daf-2* (e1370), and CF1038 *daf-16* (mu86) were maintained on Nematode Growth Medium (NGM) agar plate at 15°C. Animals at the developmental stage after the last molting and before bearing eggs (“young adult stage”) were picked up and cultured at 24°C for one day, and then transferred to a microfluidic device maintained at 25°C.

### Fabrication of microfluidic device and culture of *C. elegans* in microfluidic device

Two microfluidic devices, WormFloII and WormFloI^6^, were fabricated by combining conventional photolithography and soft lithography methods^37^. For WormFloII, a polydimethylsiloxane (PDMS) chip and a bottom PDMS plate with 1-mm thickness were assembled using the oxygen plasma bonding method. Food bacteria suspension (*E. coli,* OP50 strain with OD = 0.1 in a buffer containing 50 mM NaCl, 15 mM K_2_HPO_4_, 96 mM KH_2_PO_4_, 0.3 mM CaCl_2_, 0.3 mM MgSO_4_, 5 μg cholesterol, 1% Tween80 (Tokyo Chemical Industry Co., Ltd., Japan)) and M9 buffer with/without 1 g/L glucose were supplied to the WormFloII and WormFloI to maintain animals for observation, respectively, at 0.4 ml/h with the Micro Ceram pump (MSP-001, Yamazen Corporation, Japan).

### Observation and quantification of *C. elegans* behavior

Animals cultured in a microfluidic device were soaked in M9 buffer in a 15-cm-diameter glass dish installed in a temperature-controlled aluminum box^6^. Animals were observed under blue-light-cut illumination by a macroscope with an apochromat objective lens (1 ×) (Z16 APO, Leica, Germany) and recorded in an H264 compressed movie^6^. Animal swimming activity was measured by counting pixels in a bitmap image, in which animal movement was determined by comparing the image at the previous time frame (Supplemental Video 1)^6^. The movie compression effect on swimming activity was corrected on the activity time series by using the moving average^6^.

### Data analysis

For DMA, the fluctuation function *F*(*s*) is obtained by using the mean square root of the detrended noise round series, defined as

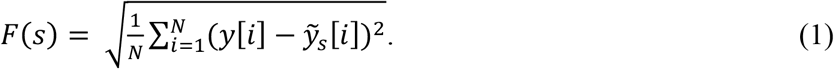

*F*(*s*) is computed from the 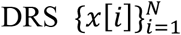 by the following procedure: the DRS {*x*[*i*]} is integrated after removing its mean value to obtain {*y*[*i*]}. This integrated DRS is filtered by the Savitzky-Golay (SG) filter to estimate the trend round series 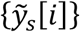, to obtain the detrended noise round series 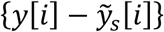. Then, the *F*(*s*) vs *s* plot on the log-log scale is fit with a linear function *y* = *ax* + *b* by the least squares method to estimate the Hurst exponent.

For DMCA, the cross-fluctuation function *F*^(1,2)^ is obtained by determining the root of cross-covariance between the bivariate detrended noise round series, defined as

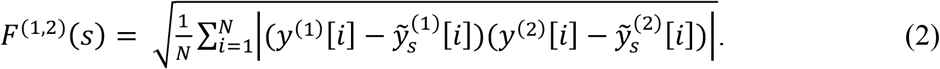

*F*^(1,2)^(*s*) is computed from bivariate DRSs 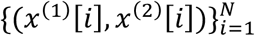 by using a procedure analogous to DMA. The linear fit to the *F*^(1,2)^ (*s*) vs *s* plot in the log-log scale can provide an estimate of the cross Hurst exponent. In addition, the multi-scale correlation coefficient *ρ*^(1,2)^(*s*), defined as

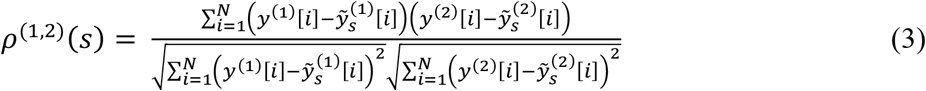

is computed to evaluate the existence of an interaction between bivariate detrended noise round series.

### Statistical analysis

Due to rejection of the normality hypothesis for scaling exponents in DMA and DMCA, and the power-law exponents of the residence-time distribution, the non-parametric Wilcoxon rank sum test was employed for pairwise comparisons between groups. In pairwise comparisons, the Benjamini and Hochberg method for correcting the false discovery rate (FDR) was used to deal with multiple testing problems.

## Supporting information

Supplementary Discussion

## Data availability

The *C. elegans* swimming activity time series and movie data reported in this paper are deposited in the Systems Science of Biological Dynamics (SSBD) database^38^, https://doi.org/10.24631/ssbd.repos.2021.11.001.

## Acknowledgement

We thank the *Caenorhabditis* Genetics Center (CGC) for *C. elegans* strains; the CGC is funded by NIH Office of Research Infrastructure Programs (P40 OD010440). We thank WormBase for planning and designing our experiments. We thank the RIKEN Center for Advanced Photonics Advanced Manufacturing Support Team for the manufacturing apparatus for culture and recording. This study was supported by JSPS KAKENHI grant number 20K20321 (Grant-in-Aid for Challenging Research (Exploratory))(Y.A.).

## Author information

### Affiliations

Cellular Informatics Laboratory, RIKEN, 2-1 Hirosawa, Wako, Saitama, 351-0198, Japan

Yusaku Ikeda, Peter Jurica, Yasushi Sako & Yukinobu Arata

Department of Mechanical Engineering, School of Engineering, Tokai University, 4-1-1 Kitakaname, Hiratsuka, Kanagawa, 259-1292, Japan

Yusaku Ikeda & Hiroshi Kimura

Graduate School of Engineering Science, Osaka University, 1-3 Machikaneyama-cho, Toyonaka, Osaka, 560-8531, Japan

Ituski Shiga & Ken Kiyono

## Contributions

Y.I., Y.A., and H.K. carried out the experiments. I.S., P.J., Y.A., and K.K. perform data analysis. K.K. and Y.S. discussed the results and commented on the manuscript. Y.A. conceived the study and wrote the manuscript.

## Corresponding author

Correspondence to Y. Arata

## Figure legends

**Fig. 3: Power-law residence-time distributions of behavioral states of fed *C. elegans***

Averaged normalized probability density distributions of active (red) and inactive states (blue) of fed wild-type (**a**, **b**), *daf-2* (**c**, **d**), and *daf-16* (**e**, **f**) animals among individual animals (grey), in log-log plot. Error bars represent standard deviations. Distributions for inactive state (**b**, **d**, **f**) were fit with linear function from −0.5 to 1.5 on x-axis (black line). Distributions for active state (**a**, **c**, **e**) were fit with linear function from −0.5 to 0.8 and were extrapolated to 1.5 on x-axis (black line).

## Extended Data Figure Legends

**Extended Data Fig. 1:**
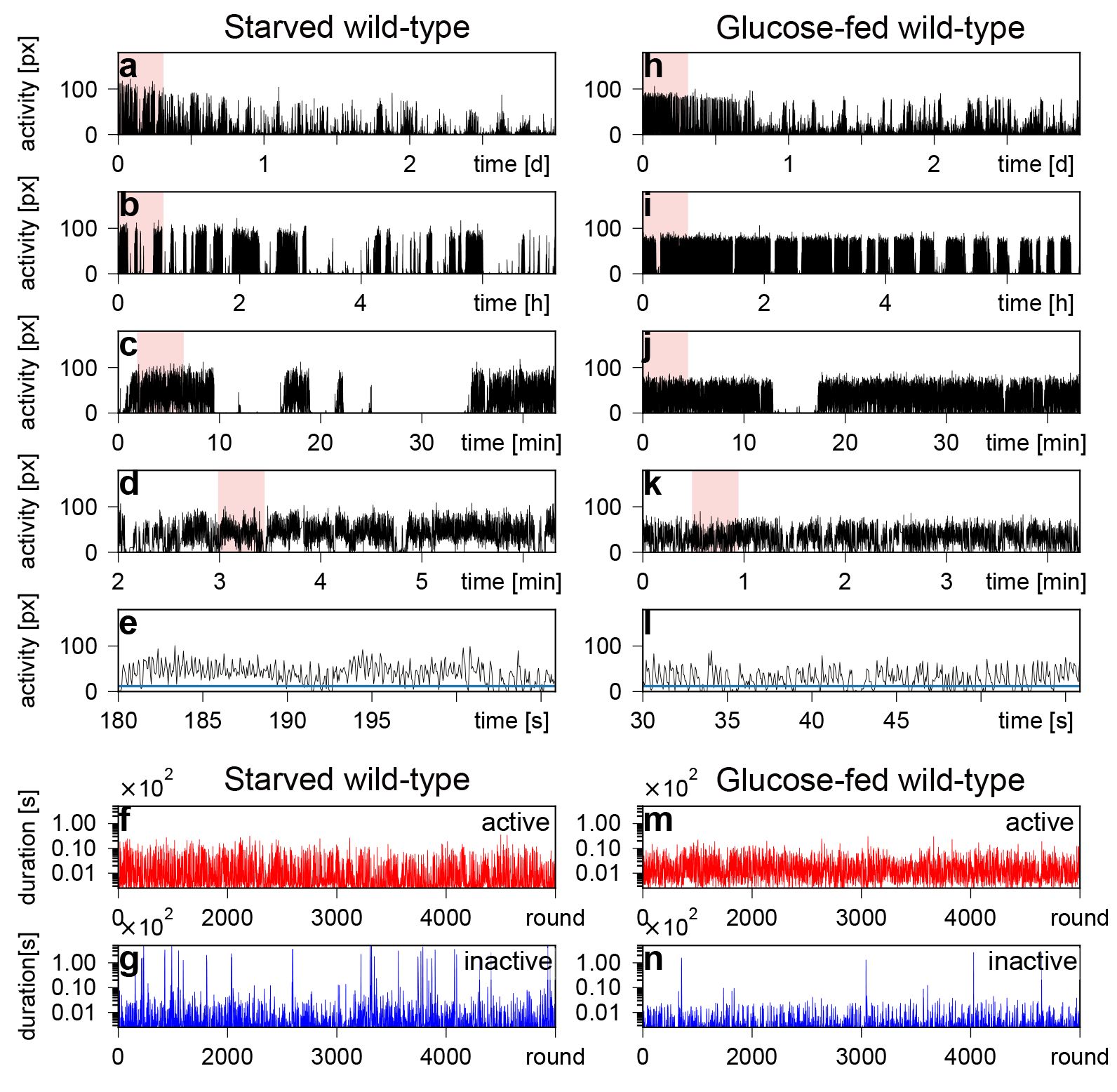
Activity time series of episodic behavior of *C. elegans* cultured without food bacteria. Swimming activity time series of starved (**a-e**) and glucose-fed (**h-l**) wild-type animals for 3 days. Red-marked regions are magnified in graphs immediately below (e.g., all of **b** represents red region from **a**; all of **c** represents red region from **b**; etc.). Active (red) and inactive (blue) DRSs in starved (**e**, **f**) and glucose-fed (**k**, **l**) wild-type animals were obtained from above activity time series for 3 days (**a**, **h**). px: number of pixels where animals moved from previous frame.

**Extended Data Fig. 2:**
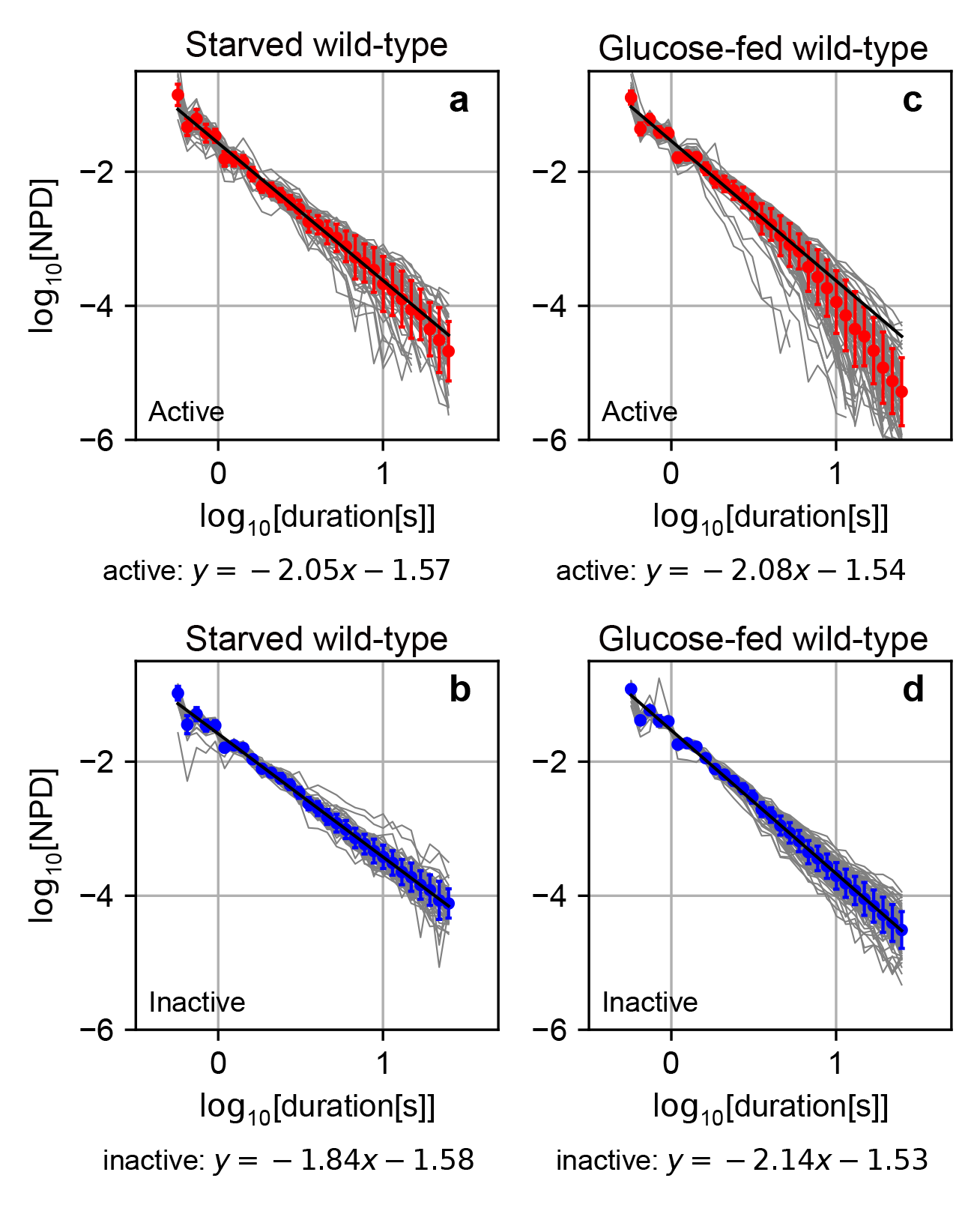
Power-law residence-time distributions of behavioral states of *C. elegans* cultured without food bacteria. Averaged normalized probability density distributions of residence time for active (red) and inactive states (blue) of starved (**a**, **b**) and glucose-fed (**c**, **d**) wild-type animals among individual animals (grey), in log-log plot. Error bars represent standard deviation. Distributions for inactive state (**b**, **d**) were fit with a linear function in a range between −0.5 and 1.5 on x-axis (black line), whereas distributions for active state (**a**, **c**) were fit with a linear function in a range between −0.5 and 0.8 and were extrapolated to 1.5 on x-axis (black line).

**Extended Data Fig. 3:**
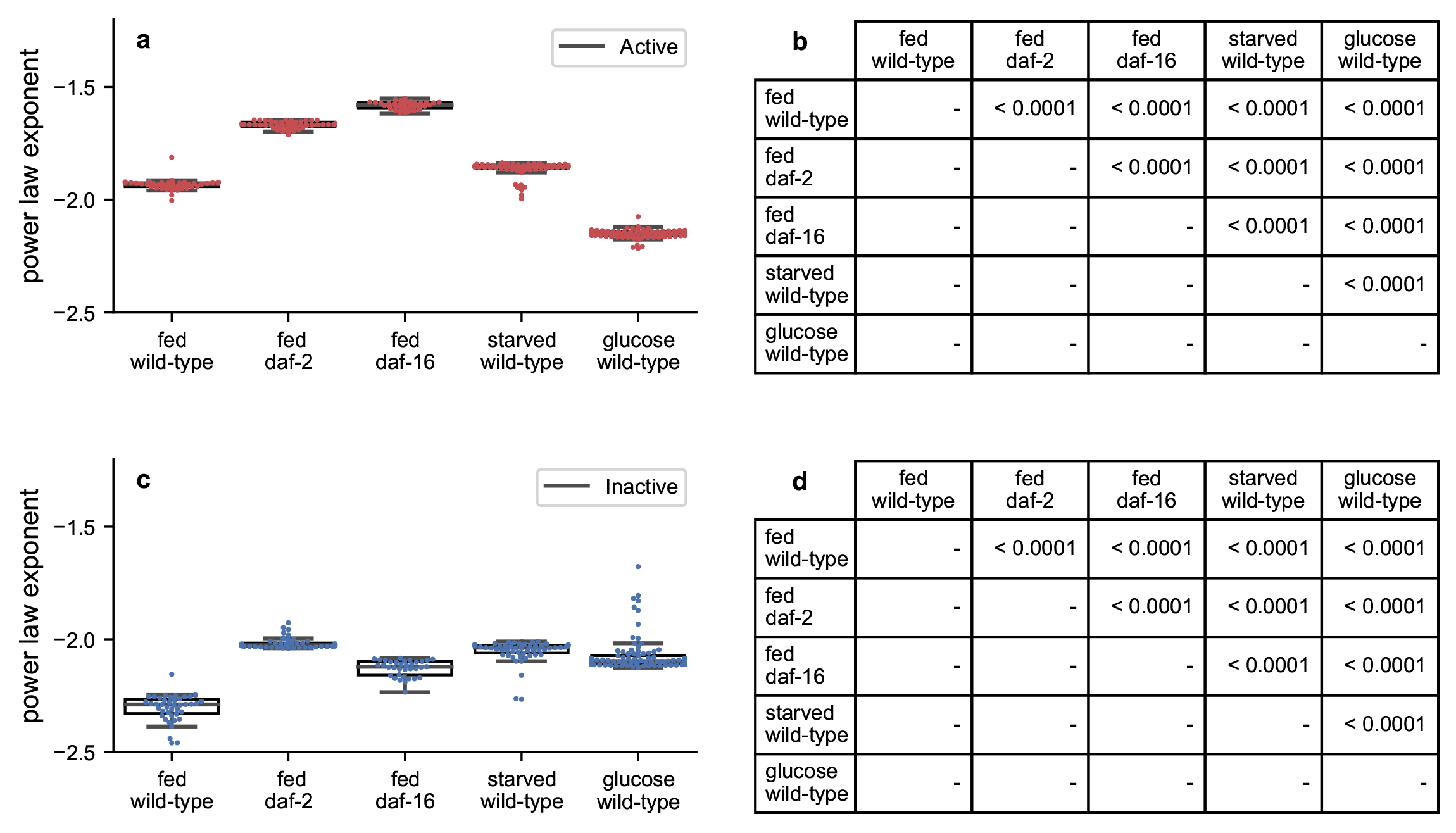
Power-law exponents of residence-time distributions for behavioral states of *C. elegans* cultured with/without food bacteria. Box-swarm plots showing raw values and medians with 25th and 75th percentiles of power-law exponents of residence times for active (**a**) and inactive (**c**) states. Error bars represent standard deviations. FDR-corrected P-values by pairwise Wilcoxon rank sum test for active (**b**) and inactive (**d**) states, with P-values > 0.05 shown in grey.

**Extended Data Fig. 4:**
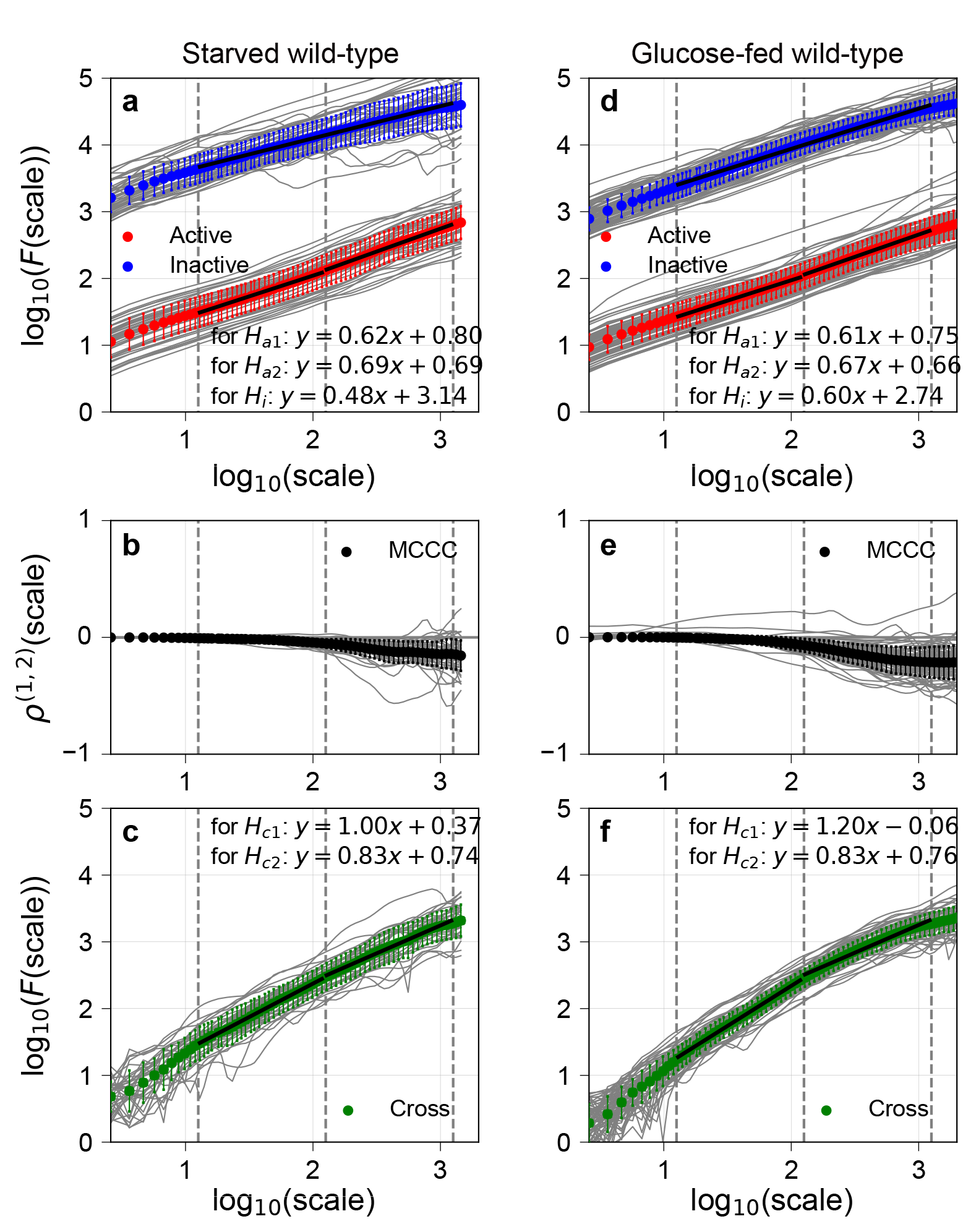
Long-range auto-/cross-correlations and multiscale cross-correlation coefficient of DRSs in fed *C. elegans*. Averaged noise function *F*(*s*) of active (red) and inactive (blue) DRSs, and averaged cross-noise function *F*^(1,2)^(*s*) between active and inactive DRSs (green), among individual animals (grey) were plotted against scale (*s*) for starved (**a**, **c**) and glucose-fed (**d**, **f**) wild-type animals. *F*(*s*) vs (*s*) plots for inactive DRS (blue) were fit with a linear function from 1.1 and 3.1. *F*(*s*) vs (*s*) plots for active DRS (red) and *F*^(1,2)^(*s*) vs (*s*) plots (green) were fit in distinct linear functions between 1.1 and 2.1 and between 2.1 and 3.1. Averaged multiscale cross-correlation coefficient between active and inactive DRSs (MCCC; black) among individual animals (grey) were plotted against scale (*s*) for starved (**b**) and glucose-fed (**e**) wild-type animals. Error bars represent standard deviations.

**Extended Data Fig. 5:**
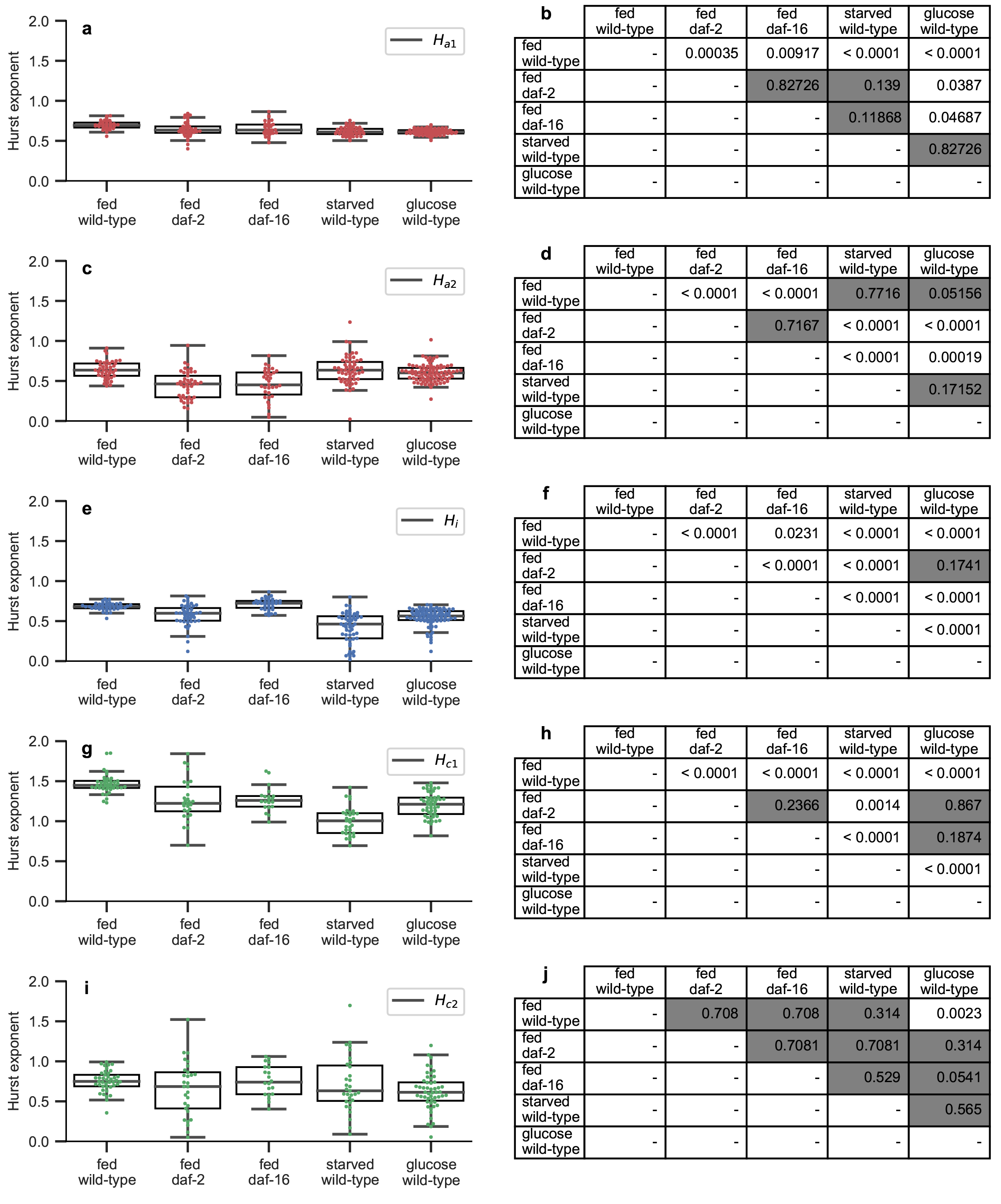
Hurst exponents of DRS in animals cultured with/without food bacteria. Raw values and medians with 25th and 75th percentiles of Hurst exponents of active DRS at shorter round scale (**a**), active DRS at longer round scale (**c**), and inactive DRS (**e**). Raw values and medians with 25th and 75th percentiles of Hurst exponents of a cross-correlated component between active and inactive DRSs at shorter round scale (**g**) and longer round scale (**i**), obtained from Fig. 4 and Extended Data Fig. 4, are shown in Box-swarm plot. FDR-corrected P-values from pairwise Wilcoxon rank sum test are shown in the corresponding combinations (**b**, **d**, **f**, **h**, **j**). P-values > 0.05 are shown in grey.

**Extended Data Fig. 6:**
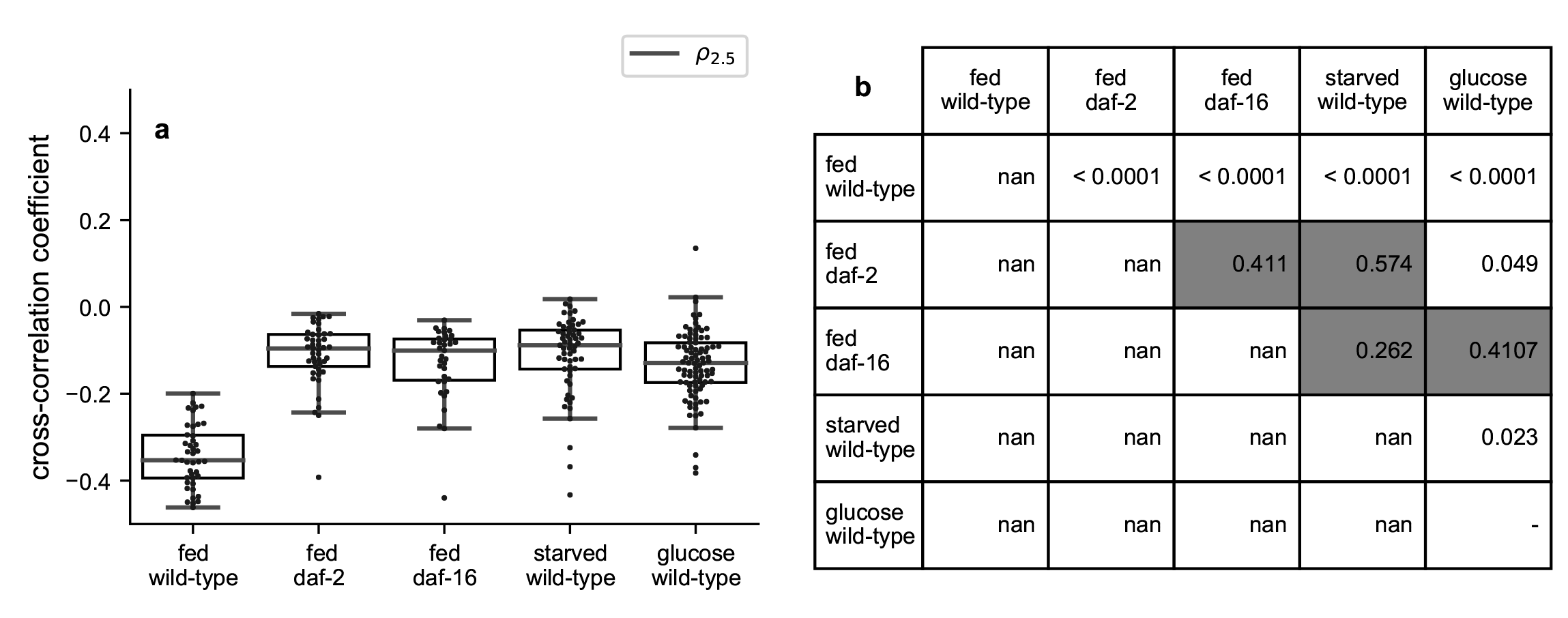
Cross-correlation coefficient between active and inactive DRSs at *log*_10_(*s*) = 2.5 in animals cultured with/without food bacteria. (**a**) Box-swarm plots showing raw values and medians with 25th and 75th percentiles of cross-correlation coefficients between active and inactive DRSs at *log*_10_(*s*) = 2.5. (**b**) FDR-corrected P-values by pairwise Wilcoxon rank sum test, with P-values > 0.05 shown in grey.

**Extended Data Fig. 7:**
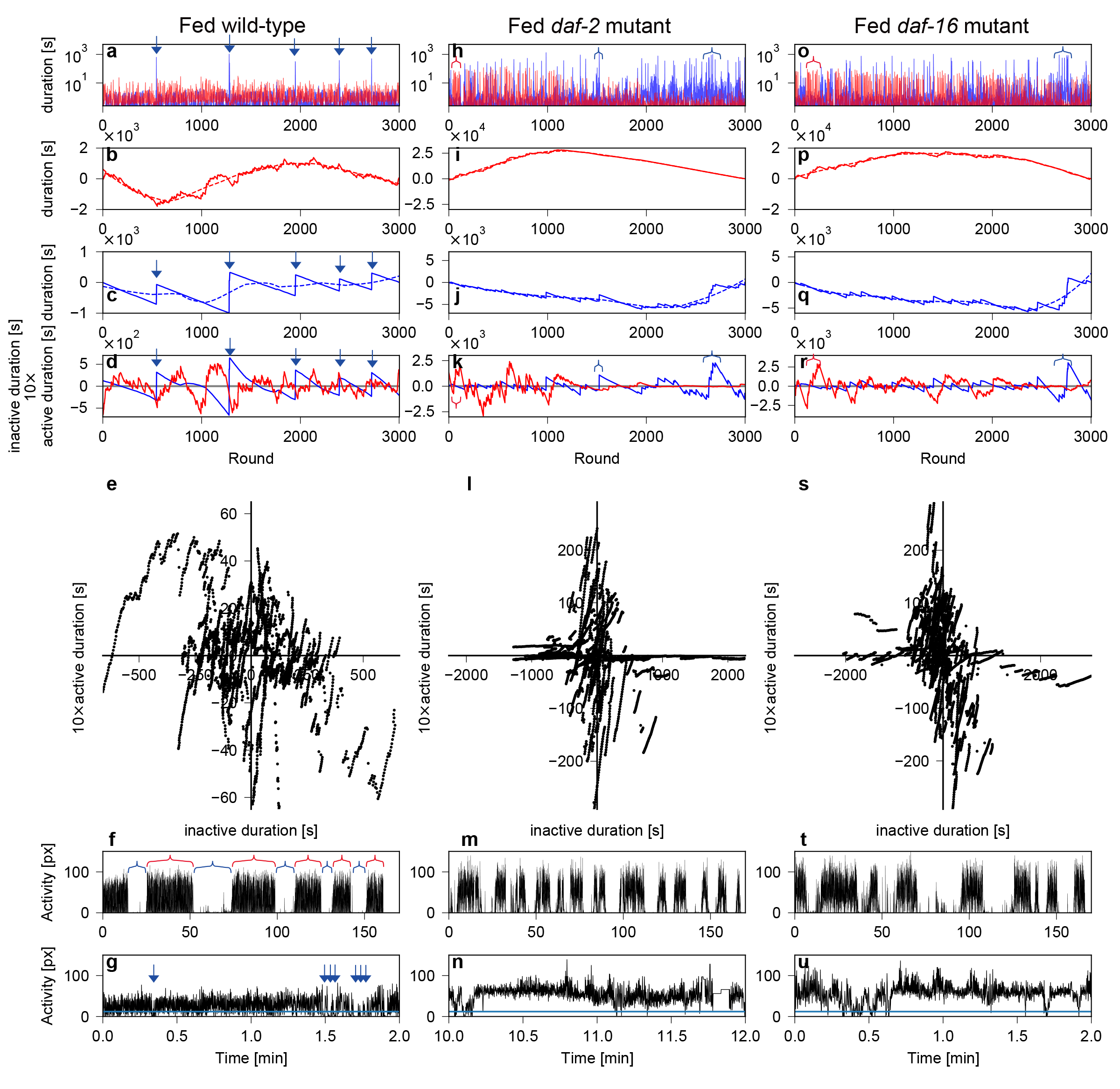
Stepwise computation of DMA and DMCA at longer round scale and the relation with activity time series. (**a**, **h**, **o**) Active (red) and inactive (blue) DRSs in fed wild-type (a), fed *daf-2* (h), and fed *daf-16* animals (o) at longer round scale (3,000 rounds). (**b-c**, **i-j**, **p-q**) Integrated DRSs, obtained by removing average durations of active (solid line; b, i, p) and inactive (solid line; c, j, q) DRSs. S-G filter was fit to integrated active and inactive DRSs, to obtain trend of integrated active (dashed line; b, i, p) and inactive (dashed line; c, j, q) DRSs. (**d**, **k**, **r**) Detrended noise round series (dNRS) for active (red) and inactive (blue) DRSs, obtained by removing trend from integrated DRSs. (**e, l, s**) Scatter plots between active dNRS (y-axis) and inactive dNRS (x-axis). (**f**) Activity time series for active and inactive DRSs in fed wild-type animals at longer round scale (3,000 rounds). (**m**, **t**) Activity time series in fed *daf-2* (m) and fed *daf-16* (t) animals, shown in the same length as fed wild-type animals (f) for comparison. (**g, n, u**) Magnification of 1/100th length of activity time series from (f, m, t). Blue arrows in (a) indicate rounds of very long inactive states in inactive DRS, which correspond to a sudden jump in integrated DRS and dNRS (blue arrows in c, d). Red and blue brackets in (h, o) indicate examples of very long active or inactive rounds that appear at high density, which correspond to a high amplitude of dNRS (red and blue brackets in k, r). Red and blue brackets in (f) indicate active and inactive episodes. Blue arrows in (g) indicate examples of short inactive states within an active episode. px: number of pixels where animals moved from previous frame.

**Extended Data Fig. 8:**
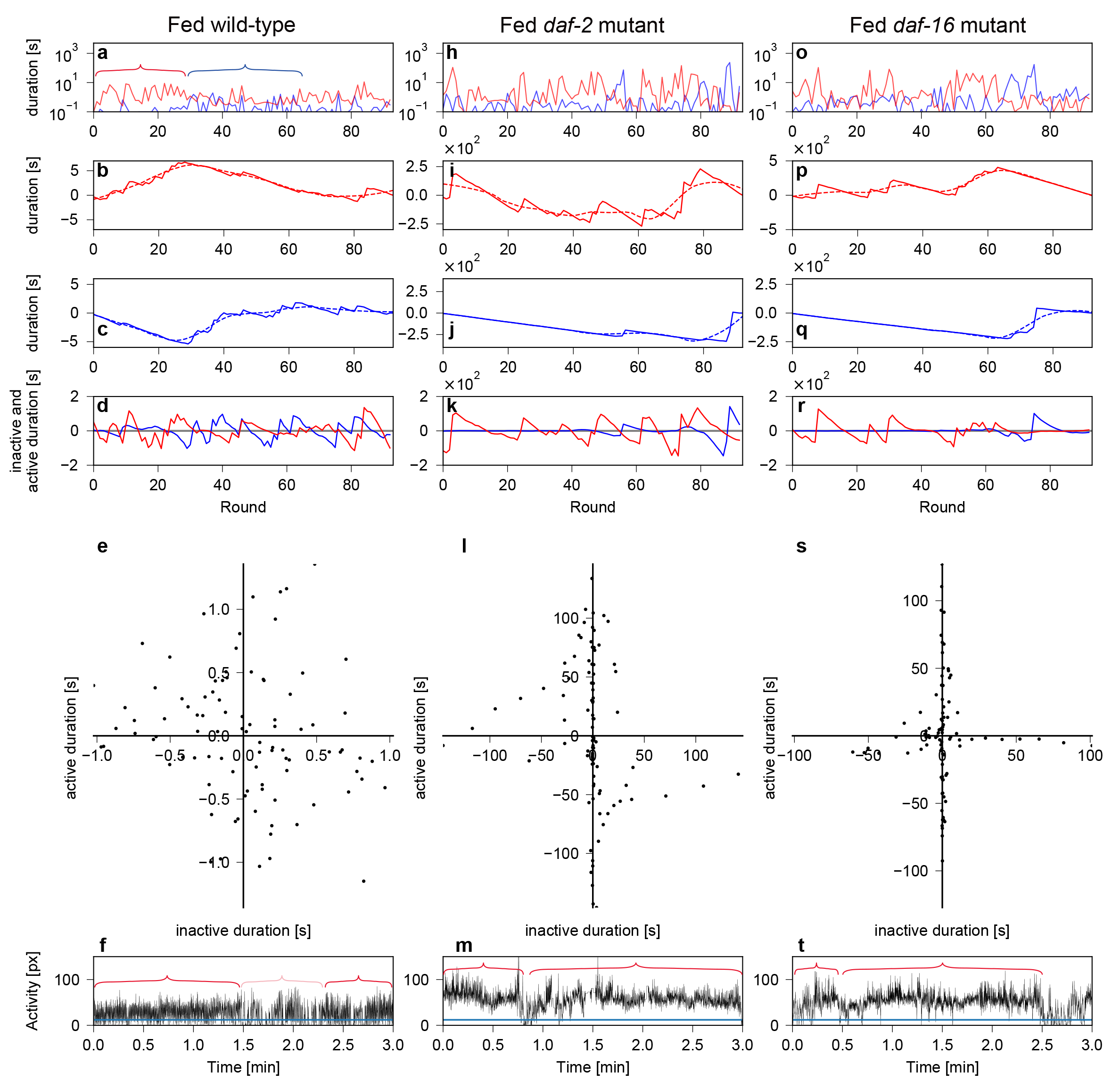
Stepwise computation of DMA and DMCA at shorter round scale and the relation with activity time series. (**a**, **h**, **o**) Active (red) and inactive (blue) DRSs in fed wild-type (a), fed *daf-2* (h), and fed *daf-16* animals (o) at shorter round scale (100 rounds). (**b-c**, **i-j**, **p-q**) Integrated DRSs, obtained by removing average durations of active (solid line; b, i, p) and inactive (solid line; c, j, q) DRSs. S-G filter was fit to integrated active and inactive DRSs, to obtain trend of integrated active (dashed line; b, i, p) and inactive (dashed line; c, j, q) DRSs. (**d**, **k**, **r**) Detrended noise round series (dNRS) for active (red) and inactive (blue) DRSs, obtained by removing the trend from the integrated DRSs. (**e, l, s**) Scatter plots between active dNRS (y-axis) and inactive dNRS (x-axis). (**f**) Activity time series for active and inactive DRSs in fed wild-type animals at shorter round scale (100 rounds). (**m, t**) Activity time series in fed *daf-2* (m) and fed *daf-16* (t) animals, shown in the same length as fed wild-type animals (f) for comparison. Red and blue brackets in DRS (a) indicate alternative appearance of consecutive rounds between longer active states/shorter inactive states (red brackets) and shorter active states/longer inactive states (blue bracket in a). Red and pink brackets in activity time series (f) indicate a time region with high swimming activity (red) or low swimming activity (pink) within a single active episode. Red brackets in (m, t) indicated active episodes. px: number of pixels where animals moved from previous frame.

## Additional Information

Supplementary Information is available for this paper.

