## Supplementary Discussion for "Fractal scaling of *C. elegans* behavior is shaped by insulin signaling"

#### Insulin signaling regulates residence-time power-law exponent in active state but not in inactive state

The absolute value of the power-law exponent in the active state of fed wild-type animals was higher than the values in fed *daf-2* and *daf-16* animals (Fig. 3a, c, e, $p < 0.05$, Extended data Fig. 3a, b) and starved wild-type animals (Extended Data Fig. 2a, $p < 0.05$, Extended Data Fig. 3a, b). Moreover, the absolute value of the power-law exponent in the active state of starved wild-type animals was restored in glucose-fed wild-type animals (Extended Data Fig. 2c, $p < 0.05$, Extended Data Fig. 3a, b). These results indicate that insulin signaling can regulate the power-law exponent for the active state. Although the absolute value of the power-law exponent of the inactive state was similarly increased in fed wild-type animals compared to fed *daf-2*, fed *daf-16* (Fig. 3b, d, f), and starved wild-type animals, the exponent of the inactive state of starved wild-type animals was not significantly restored in glucose-fed wild-type animals (Extended Data Fig. 2b, d, Extended Data Fig. 3c, d). Therefore, we conclude that the power-law exponent of the active state is regulated by insulin signaling.

#### Experimental results to test insulin signaling-dependent control of weak fractal memories in active DRS at long-round scale and weak fractal memories in inactive DRS at the whole round scale

$H_{a2}$ and $H_i$ were not systematically or significantly affected by the interventions for insulin signaling ON (fed and glucose-fed wild-type) and OFF (fed *daf-2* and fed *daf-16*, and starved wild-type) situations. Compared to the $H_{a2}$ in fed wild-type animals ($H_{a2} = 0.72$ in Fig. 4a), the $H_{a2}$ was lower in insulin-signaling mutants ($H_{a2} = 0.49$ in either fed *daf-2* or fed *daf-16* animals, Fig. 4d, g, $p < 0.05$ in Extended Data Fig. 5c, d), consistent with the insulin signaling-dependent control of $H_{a2}$. However, $H_{a2}$ was not significantly reduced in starved wild-type animals ($H_{a2} = 0.69$) and was not restored in glucose-fed wild-type animals ($H_{a2} = 0.67$, Extended Data Fig. 4a, b, Extended Data Fig. 5c, d). Compared to the $H_i$ in fed wild-type animals ($H_i = 0.68$ in Fig. 4a), $H_i$ was decreased in two insulin-signaling OFF situations ($H_i = 0.60$ in fed *daf-2* animals in Fig. 4d and $H_i = 0.48$ in starved wild-type animals in Extended Data Fig. 4a, respectively, $p < 0.05$ in Extended Data Fig. 5e, f). $H_i$ in starved wild-type animals ($H_i = 0.48$) was restored in glucose-fed wild-type animals ($H_i = 0.60$ in Extended Data Fig. 4a, d, $p < 0.05$ in Extended Data Fig. 5e, f). These results are consistent with the insulin signaling-dependent control of $H_i$. However, $H_i$ in fed wild-type animals ($H_i = 0.68$

in Fig. 4a) was not reduced in fed *daf-16* animals ($H_i = 0.71$, respectively in Fig. 4g). Together, these results do not provide strong evidence of the insulin signaling-dependent control of $H_{a2}$ and $H_i$.

**Insulin signaling-dependent control of temporal behavioral activity patterns**

Our DRS-based fractal analyses showed that insulin signaling modulates the mechanism for determining the residence-time distribution in fractal kinetics A1 and A2 (Fig. 3), and the mechanism for determining fractal memory in fractal kinetics C ($H_{c1}$; Fig. 4a, d, g). Insulin signaling also modulates the switch for the lateral linking mechanism between fractal kinetics A2 and I ($\rho^{(1,2)}(s)$; Fig. 4) to shape fractal scaling of *C. elegans* behaviors (Fig. 5). With the activation of insulin signaling, the power-law residence-time distribution in the active state changes to an exponential-like distribution at the longer time scale, the fractal memory in fractal kinetics C is strengthened, and the lateral linking mechanism between fractal kinetics A2 and I is activated. To understand how the insulin signaling-dependent behavioral control detected by DRS-based analyses altered the temporal activity patterns of *C. elegans* behavior, we studied the relation between the DRS and the behavioral activity time series.

First, we investigated the insulin signaling-dependent control of the larger absolute power-law exponent at the shorter time scale and the exponential-like decline at longer time scale in the residence-time distribution of the active state (Fig. 3). The larger absolute power-law exponent and exponential-like decline were both caused by a relative decrease of the appearance frequency of longer active states to shorter active states at each of the shorter and longer time scales. In the activity time series, we noticed that active episodes have a characteristic temporal pattern, wherein a series of active states was generated by frequent interruptions by short inactive states in fed wild-type animals, which was caused by a series of short active rounds in active DRS (Fig. 2d, e, Extended Data Fig. 7f, g). In active episodes in fed *daf-2* and fed *daf-16* mutant animals, the insertion of the short inactive states in an active episode was significantly reduced (Fig. 2k, l, r, s, Extended Data Fig. 7n, u), resulting in more frequent long active states in fed *daf-2* and fed *daf-16* animals (compare Fig. 2f to Fig. 2m, t). Therefore, one cause of the larger absolute power-law exponent and exponential-like decline was the characteristic temporal pattern of active episodes in fed wild-type animals.

To identify how animals behaved during the frequent short inactive states, we observed swimming motions of fed wild-type animals from movies. We found that the short inactive states were derived from a characteristic posture, wherein fed wild-type animals transiently kept a body-bent posture at a sub-second time scale after a continuous alternative bent posture for swimming (Supplementary Video 1), which we previously reported as the characteristic "posing" posture[1]. In the movies of fed *daf-2* and fed *daf-16* mutants, animals took the posing posture less frequently (Supplementary Video 2, 3). Although the ethological role and physiological mechanisms of posing remain unknown, the frequency of posing is under the control of insulin signaling.

Second, we investigated insulin signaling-dependent control of the negative correlation coefficient between the active and inactive DRSs at the long-round scale (3,000 rounds, $\rho_{2.5}$, Fig. 4) as evidence for an ON/OFF switch on a lateral linking mechanism between fractal kinetics A2 and I. To seek the cause, we computed a multiscale cross-correlation coefficient by focusing on the inactive DRS in fed wild-type animals as follows. In step 1, the inactive DRS (Extended Data Fig. 7a) was integrated after removing the average of residence times in the DRS (Extended Data Fig. 7c). The integrated inactive DRS showed several rounds of sudden large jumps (blue downward arrows, Extended Data Fig. 7c), which were derived from very long inactive states in the original DRS (blue downward arrows, Extended Data Fig. 7a). After the sudden large jumps in the integrated inactive DRS, the DRS values continuously decreased (Extended Data Fig. 7c). This continuous decreasing corresponded to a series of short inactive states in the original DRS at the interval between two very long inactive states (Extended Data Fig. 7a).

In step 2, from the integrated inactive DRS, we obtained the trend round series by applying the S-G filter (dashed line, Extended Data Fig. 7c), and subsequently obtained the detrended noise round series (dNRS) (Extended Data Fig.7d). Sharp peaks of noise amplitude in the inactive dNRS were derived from the very long inactive states in the original inactive DRS. Similarly, the relation between dNRS and DRS can be found in the active DRS (Extended Data Fig. 7a, b, d). Inactive dNRS and active dNRS were in an antiphasic trend (Extended Data Fig.7d), wherein a positive (negative) amplitude of inactive dNRS tended to be associated with a negative (positive) amplitude of inactive dNRS. This antiphasic trend caused the negative cross-correlation at the long-round scale detected by multiscale correlation coefficient (Fig. 4b). To reveal the relation of this antiphasic trend in dNRS with the activity time series, we focused on the sudden jumps and subsequent constant decrease in inactive dNRS and DRS (Extended Data Fig.7a, d). Very long inactive rounds in the original inactive DRS corresponded to an inactive episode in activity time series (blue brackets, Extended Data Fig.7f). On the other hand, a series of short inactive states after very long inactive states in inactive DRS corresponded to frequent insertion of short inactive states in an active episode in the activity time series (red brackets and blue arrows, Extended Data Fig.7f, g, respectively). Based on this understanding, we concluded that the negative correlation between active and inactive DRSs in fed wild-type animals was derived from a characteristic alternative switching between active and inactive episodes and internal structure in active episode.

In fed *daf-2* and *daf-16* animals, we did not find obvious defects in the antiphasic trend in the active and inactive dNRSs (Extended Data Fig. 7k, r). To identify the cause of the shrunken negative correlation in fed mutant animals (Fig. 4e, h), we made a scatter plot between the active and inactive dNRSs. The scatter plot for fed wild-type animals showed a weak downward trend centered at the origin (Extended Data Fig. 7e), which was consistent with a weak negative correlation ($\rho_{2.5} = -0.4$,

Fig. 4b). In fed *daf-2* and fed *daf-16* animals, the scatter plots distributed along the axes in a cross-shaped manner, suggesting that one of noise amplitudes in the active or inactive dNRS was randomly enlarged (Extended Data Fig. 7l, s). We confirmed this random enlargement (Extended Data Fig.7k, r; note that the vertical axis of dNRS is 100× larger than that of fed wild-type animals in Extended Data Fig. 7d). We conclude that the random enlargement of noise amplitude in active or inactive dNRS caused the shrunken cross-correlation coefficient in fed mutant animals.

The random noise amplitude enlargement in the active or inactive dNRS was caused by the consecutive rounds of very long active or inactive states in the DRSs (compare red and blue brackets for active and inactive rounds in Extended Data Fig. 7k, r and Extended Data Fig. 7h, o). Such consecutive very long active or inactive states were not observed in fed wild-type animals (Extended Data Fig. 7a). These disordered structures in active and inactive DRSs in fed mutant animals were likely derived from disrupted patterns of alternative switching of active and inactive episodes in the activity time series (Extended Data Fig. 7m, t) and the disrupted internal structure of an active episode in the activity time series (Extended Data Fig. 7n, u). Together, these data indicate that the negative correlation coefficient between the active and inactive DRSs at the *long*-round scale was derived from the characteristic alternative switching of active and inactive episodes and the internal structure of an active episode in the activity time series in fed wild-type animals.

Third, we investigated insulin signaling-dependent control of strong fractal memory in a pseudo cross-correlated component between active and inactive DRSs at the *short*-round scale (100 rounds, $H_{c1} = 1.47$, Fig. 4a). To identify the cause of strong fractal memory, we tracked the process of DMCA, which is basically the same as the process to compute a multiscale cross-correlation coefficient. In active and inactive DRSs, we found that some fed wild-type animals displayed a consecutive series of rounds with relatively longer active and shorter inactive states, followed by a consecutive series of rounds with relatively shorter active and longer inactive states (red and blue brackets, Extended Data Fig. 8a). Each of these consecutive rounds corresponded to gradual increase of integrated active and inactive DRSs (Extended Data Fig. 8b, c). During a gradual increase, relatively large positive noises tended to appear in the dNRS (Extended Data Fig. 8d). Consequently, the noise amplitudes of the active and inactive states tended to change in an antiphasic manner (red and blue brackets, Extended Data Fig. 8d). A negative correlation was also observed in the scatter plot, although this correlation was very weak (Extended Data Fig. 8e). This weak negative correlation was mathematically incorporated to compute $F^{(1,2)}(s)$ in DMCA (see equation (2)). In this way, we considered that this weak negative correlation between active and inactive dNRSs may be, at least partially, a cause of the strong fractal memory. The alternative appearance of consecutive rounds between longer active/shorter inactive states and shorter active/longer inactive states in DRSs was derived from another characteristic internal structure in an active episode in the activity time series.

Namely, a high swimming activity-prone time region and a low activity-prone time region alternatively appeared (red and pink brackets, Extended Data Fig. 8f). Strong fractal memory detected by DMCA may be derived from a scale-dependent change of such internal structure of an active episode.

In fed *daf-2* and *daf-16* animals, we did not find obvious defects in the antiphasic pattern with weak negative correlation between the active and inactive dNRSs at the short-round scale (Extended Data Fig. 8k, r). Scatter plots for fed mutant animals showed that dNRS values tended to distribute along the axes in a cross-shaped manner (Extended Data Fig. 8l, s). In the dNRS of fed mutant animals, one of noise amplitudes in the active or inactive dNRS was randomly enlarged (Extended Data Fig. 8k, r; vertical axis of the dNRS was 100× larger than that of fed wild-type animals, Extended Data Fig. 8d). Random enlargement of the noise amplitude in the active or inactive dNRS disrupted the negative correlation in the short-round scale, which was mathematically incorporated to compute $F^{(1,2)}(s)$ in DMCA (see equation (2)). In this way, random enlargement of the noise amplitude in active or inactive dNRS may be involved in the weakened strong fractal memory in fed mutant animals. The alternative appearance of high/low activity-prone time regions disappeared within an active episode in the activity time series in fed *daf-2* and fed *daf-16* animals (red brackets, Extended Data Fig. 8m, t). Therefore, we conclude that fractal scaling of a characteristic internal structure within active episodes in fed wild-type animals is an insulin-dependent behavior.

In summary, by investigating the relation between DRS and activity time series, we found that two types of characteristic internal temporal patterns in active episodes and alternative switching patterns between active and inactive episodes are insulin signaling-dependent behavioral patterns in *C. elegans*.

### Supplementary Methods

#### Design, fabrication, and characterization of microfluidic device

For culture chambers and the junctional micro-slits in WormFloII, which have different heights, a mold master for the PDMS chip was fabricated by two-step patterning of a negative-type photoresist (SU-8 2000 series, MicroChem, USA). A PDMS prepolymer made with a 10:1 mass ratio of PDMS (dimethylpolysiloxane, Silpot 184, Dow Corning Toray) and a curing reagent were spread on mold masters by a spin coater to control the thickness at 0.4 mm. The PDMS prepolymer on the mold masters was cured at 75°C for 2 h. After solidifying the PDMS prepolymer, a 0.1-mm diameter hole was made by $CO_2$ laser at the center of a culture chamber for animal loading (Epilog Zing, USA). Inlet and outlet ports were made by a trepan puncher (⌀ 2 mm, Kai, Japan).

We tested the microfluidic device for uniform flow across the chambers. Briefly, M9 buffer (containing red food coloring) was supplied into the device with Micro Ceram pump (MSP-001,

Yamazen Corporation, Japan) at the same speed (0.4 ml/h) as for culturing *C. elegans*. Uniform distribution of red food coloring among all chambers was tested (Supplementary Video 4).

**Culture of *C. elegans* in microfluidic device**

Tubes were attached to inlet and outlet ports of a microfluidic device, which was fixed in a glass dish (15-cm diameter and 5.4-cm depth) with an apparatus made of polyether ether ketone (PEEK). Next, the microfluidic device was soaked with M9 buffer. The microfluidic device in the glass dish was sterilized in an autoclave at 121°C for 15 min. *C. elegans* animals were installed into the sterilized microfluidic device by using a glass pipette from a hole on the roof of each chamber in the device. After installing animals, holes on the device were sealed with an ethanol-sterilized PDMF sheet (0.1-mm thickness), which was fixed with an ethanol-sterilized acrylic board that was connected to the basal PEEK apparatus.

For observation, the microfluidic device in the glass dish was inserted into a temperature-controlled aluminum box under a macroscope with an apochromat objective lens (1 ×) (Z16 APO, Leica, Germany)[1]. To maintain the quality of the OP50 suspended medium during recording, the medium was stored at 4°C and was continuously supplied to the microfluidic device via 3-meter-long sterilized polyvinyl chloride tubes (⌀ 1 mm, Tygon LMT-55, Saint-Gobain, France) held at room temperature. A 3-meter-long sterilized tube was connected to a drip chamber used for intravenous transfusion (ISA-200A 00 Z, NIPRO, Japan) located at 10-cm before the device inlet. The drip chamber removed air bubbles that appeared in the tube due to the temperature increase from 4°C. Medium was perfused at 0.4 ml/h flow through a Micro Ceram pump (MSP-001, Yamazen Corporation, Japan), which was located at the tube downstream from the device outlet. Outlet flow was discarded to a waste bottle. Illumination light and temperature controls at the culture chambers on the device in the recording system were qualified as described in our previous study[1].

### Supplementary Figure Legends

**Supplementary Video 1: Fed wild-type animals cultured with food bacteria**

**(Upper panel)** *C. elegans* swimming behavior in fed wild-type animals. Pixels where animals moved from previous frame are shown in green. **(Lower panel)** Swimming activity (number of green pixels within culture chamber) of animal ID 26 (indicated by red frame in above movie) is highlighted by green line. Vertical dashed line indicates time point shown in frame in above movie. Blue lines at high (active state) and low (inactive state) values indicate activities above or below activity threshold of 12 pixels/frame, respectively. Posing was observed at 8.5, 10, 19.5, 22.5, 15.5, 28.5, 40, 43.5, and 50 sec.

**Supplementary Video 2: Fed *daf-2* animals cultured with food bacteria**

**(Upper panel)** *C. elegans* swimming behavior in fed *daf-2* animals. Pixels where animals moved from previous frame are shown in green. **(Lower panel)** Swimming activity (number of green pixels within culture chamber) of animal ID 27 (indicated by red frame in above movie) is highlighted by green line. Vertical dashed line indicates time point shown in frame in above movie. Blue lines at high (active state) and low (inactive state) values indicate activities above or below activity threshold of 12 pixels/frame, respectively. Posing was observed at 57 and 58.5 sec.

**Supplementary Video 3: Fed *daf-16* animals cultured with food bacteria**

**(Upper panel)** *C. elegans* swimming behavior in fed *daf-16* animals. Pixels where animals moved from previous frame are shown in green. **(Lower panel)** Swimming activity (number of green pixels within culture chamber) of animal ID 23 (indicated by red frame in above movie) is highlighted by green line. Vertical dashed line indicates time point shown in frame in above movie. Blue lines at high (active state) and low (inactive state) values indicate activities above or below activity threshold of 12 pixels/frame, respectively. Posing was observed at 8, 15, and 48.5 sec.

**Supplementary Video 4: Uniform flow among chambers on WormFloII**

M9 buffer containing red food coloring was supplied to the microfluidic device. The weak grey signal that starts to appear in the flow channel at 3 sec is the signal from the red food coloring.
